# Phosphorylation of a tumor-derived ASXL2 epitope remodels peptide-HLA binding affinity and interaction dynamics

**DOI:** 10.64898/2026.03.08.710418

**Authors:** Jiahui Zhang, Lin Lv, Bairun Chen, Xinpei Yi

**Author notes:** These authors contributed equally (Jiahui Zhang, Lin Lv). Correspondence (X.Y.).

## Abstract

The expansion of cancer immunotherapy has made it increasingly important to identify tumor-specific HLA class I epitopes and to define the molecular determinants of peptide–HLA binding and stability. Post-translationally modified epitopes, particularly phosphorylation, can reflect oncogenic signaling states and offer highly tumor-selective targets. Despite their potential, the structural and dynamic mechanisms by which phosphorylation reshapes peptide–HLA affinity and stability remain unclear. Here, we investigate a phosphorylated ASXL2-derived epitope supported by confident MS/MS-based phosphosite localization and detected across multiple cancer types. We performed all-atom molecular dynamics simulations of peptide–HLA (pHLA) complexes to compare the phosphorylated peptide, its non-phosphorylated counterpart, and related variants. Our simulations indicate that phosphorylation reconfigures non-bonded interaction networks within the binding groove by introducing new local contacts, leading to an enhanced predicted binding affinity. Furthermore, principal component analysis revealed that phosphorylation increases the overall conformational flexibility and alters the collective backbone motions of the pHLA complex. Together, these findings provide a structural–dynamic basis for how phosphorylation can modulate pHLA stability and interaction dynamics, guiding the rational prioritization and targeting of cancer-specific pHLA complexes for immunotherapy.

## Introduction

The presentation of antigenic peptides by human leukocyte antigen (HLA) molecules is the crucial first step in triggering adaptive immune recognition [1,2]. Intracellularly, proteins are continuously processed into short peptides [3], a specific subset of which is loaded into the binding groove of HLA class I molecules and presented on the cell surface as peptide–HLA (pHLA) complexes [4]. Circulating T cells continuously survey these pHLA complexes via their T cell receptors (TCRs) [5]. Recognition of a non-self or aberrant peptide triggers robust T-cell activation and downstream effector responses aimed at eliminating the presenting cell, a mechanism central to tumor clearance [6]. Consequently, deciphering the exact properties of peptide–HLA interactions is paramount to cancer immunotherapy [7,8], serving as the structural blueprint for the rational design of therapeutic vaccines [9], engineered TCRs [10], and TCR-mimic antibodies [11].

Beyond canonical targets, post-translationally modified (PTM) antigenic peptides introduce a biologically informative layer to the pHLA landscape. In malignancies, aberrant signaling and metabolism produce a unique repertoire of modified peptides—most notably through phosphorylation, acetylation, and methylation. These PTM neoantigens represent a highly significant class of therapeutic targets: they not only offer enhanced tumor selectivity but can also be shared across diverse patient populations, eliciting robust immunogenicity [12]. Despite their immense clinical potential, these modified antigens have been largely overlooked by previous studies. This historical neglect stems from a strict technical bottleneck: because conventional genomic and transcriptomic approaches cannot capture these dynamic post-translational events, PTM antigens can exclusively be detected through proteomics-level mass spectrometry. Today, tandem mass spectrometry (MS/MS) within immunopeptidomics has become indispensable, providing the only direct, site-resolved evidence for PTM identity and localization [13], thereby enabling the confident discovery of these elusive targets.

Yet, successful mass spectrometry identification solves only half of the equation; interpreting the therapeutic potential of these PTM neoantigens requires a deep understanding of their binding biophysics [14]. Recently, some researchers have begun to investigate how modifications like phosphorylation affect antigen–HLA interactions, developing tools to assess the binding affinity and static structures of phosphorylated pHLA complexes [15–17]. However, these investigations are strictly confined to static snapshots. Furthermore, the vast majority of current predictive models and tools remain optimized almost exclusively for unmodified canonical sequences [18,19]. Consequently, the precise biophysical mechanisms by which a phosphorylation event dynamically remodels the peptide–HLA interface and alters overall complex stability remain completely unresolved. Because pHLA dynamics are crucial for governing T-cell recognition but are notoriously difficult to probe experimentally in vitro [20], in silico molecular dynamics (MD) simulations are essential to sample conformational fluctuations over time [21]. While MD methods have been widely applied to study standard unmodified pHLA systems [22–27], comprehensive dynamical characterizations of phosphorylated complexes remain remarkably limited.

To address this critical gap, we focus on a specific phosphorylated antigenic peptide, KVIpSPSQKHSK, from ASXL transcriptional regulator 2 (ASXL2) (**Figure 1A, top**; **Figure S1**) [28]. ASXL2 is an epigenetic regulator with multiple functions [29]. Moreover, emerging evidence underscores its clinical significance in various malignancies, where its aberrant expression is associated with tumorigenesis and poor prognosis in colorectal and pancreatic cancers, as well as immunomodulation in head and neck squamous cell carcinoma [30–32]. Interestingly, Figure 1A (bottom) shows that this peptide was exclusively detected in 7 tumor samples in the caAtlas database [33] but was completely absent in normal tissues. This tumor-specific distribution spans multiple cancer types, including melanoma, leukemia, meningioma, and ovarian cancer, as evidenced by MS/MS data. Preliminary pan-cancer analysis leveraging the CPTAC dataset [34,35] reveals that ASXL2 is significantly dysregulated across multiple omics layers, including mRNA expression, protein abundance, and site-specific phosphorylation (**Figures 1B-D**). These consistent alterations between tumor and normal tissues underscore the potential clinical relevance of this antigen and its post-translational modifications. Furthermore, our pan-cancer pathway analysis reveals a distinct pattern where this phosphorylation correlates with upregulated tumorigenic signatures and dampened immune surveillance pathways (**Figure 1E**). These data point to a potential oncogenic mechanism where the phospho-peptide fosters an immunosuppressive microenvironment, hindering effective anti-tumor immunity.

**Figure 1.**
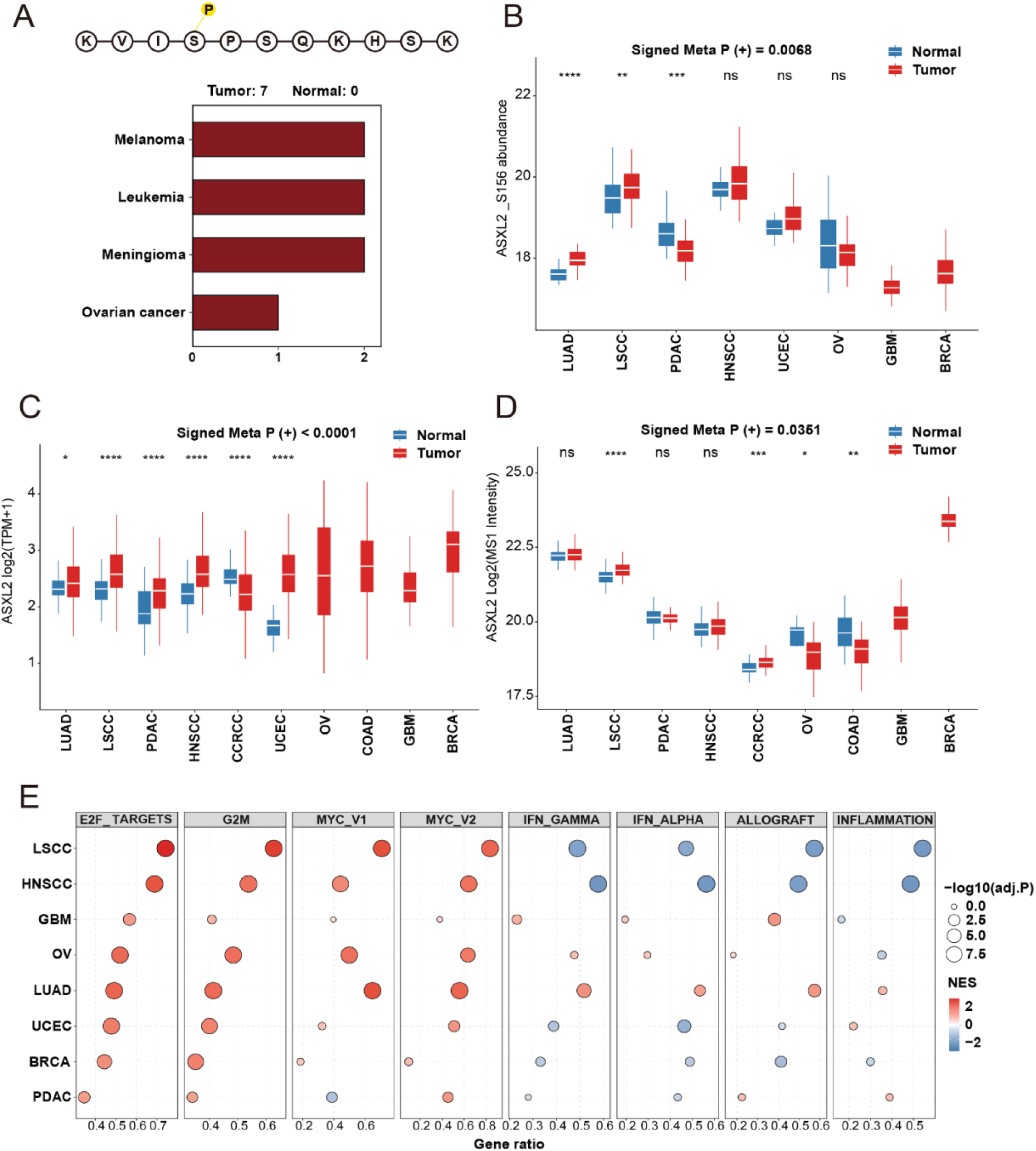
Identification and multi-omics characterization of the tumor-specific ASXL2 phospho-peptide. (A) Schematic sequence (top) and detection frequency statistics in the caAtlas database (bottom) of the ASXL2 phospho-peptide. (B–D) Pan-cancer dysregulation of ASXL2 across multiple omics layers. Boxplots compare the abundance of (B) the specific S156 phosphosite, (C) mRNA expression, and (D) total protein levels between tumor (red) and normal (blue) tissues in the CPTAC dataset. The Signed Meta P-value indicates the global statistical significance of the upregulation in tumors. (E) Functional characterization of ASXL2 phosphorylation via Gene Set Enrichment Analysis (GSEA).

In this work, we performed all-atom molecular dynamics (MD) simulations of the MS/MS-validated ASXL2-derived peptide bound to HLA-A*31:01, as described in Materials and Methods section 5, and obtained trajectories for the phosphorylated, non-phosphorylated, and related pHLA systems. We then analyzed non-bonded interactions and found substantial differences between the phosphorylated and non-phosphorylated complexes. Free-energy calculations further indicated that phosphorylation markedly alters binding affinity. To compare the conformational dynamics, we performed principal component analysis (PCA), which showed that phosphorylation strongly affects backbone motions within the pHLA complex. Together, these results provide a detailed structural and dynamical view of how phosphorylation reshapes a clinically relevant antigenic peptide–HLA complex, and they establish a framework for studying PTM antigen presentation and pHLA-targeted cancer immunotherapies.

## Results

### Benchmarking the force field and assessing trajectory stability

We first evaluated whether the chosen force field can distinguish between phosphorylated peptide–HLA systems with binding versus non-binding behavior. To this end, we assembled 50 binding and 50 non-binding phosphorylated pHLA systems as described in the Methods section. In principle, sufficiently long simulations could allow peptides from non-binding systems to dissociate from HLA; however, direct dissociation is rarely observed within practical computational budgets. We therefore performed short MD simulations and compared the relative stability of the binding and non-binding sets. As one stability proxy, we quantified peptide residue flexibility using the root-mean-square fluctuation (RMSF) [36]. To reduce trapping in local energy minima, simulations were conducted at elevated temperature [37]. Notably, related experimental approaches also infer binding differences from stability measurements [38]. For consistency, we selected all phosphorylated peptides to a length of nine residues, the most common length for HLA class I ligands [33] and compared per-residue RMSF values (**supplementary Figure 2**). The median RMSF values of the binding set were lower than those of the non-binding set, most prominently at positions 1, 2, 8, and 9; several positions showed statistically significant differences (positions 5, 6, 8, and 9). Together, these results support the suitability of the chosen force field for differentiating phosphorylated peptide–HLA binding from a statistical perspective. The peptide sequences and corresponding HLA alleles are provided as Supplementary file 1.

To verify the robustness of our simulation protocol, we report the RMSD traces for all production runs (**supplementary Figure 3**) and RMSF profiles for the phosphorylated, non-phosphorylated, and protonated phosphorylated systems (**supplementary Figure 4**). Across all simulations, RMSD values remained within a narrow range, indicating that the systems were stable over the production window. The RMSF profiles showed elevated flexibility primarily at protein termini whereas non-terminal residues remained stable, which is expected. Together, these analyses support that our simulation protocol is robust and suitable for the subsequent analyses.

### Phosphorylation alters non-bonded interactions locally and globally

The local environments around the focal site in the phosphorylated and non-phosphorylated systems are shown in Figure 2. In the enlarged views, the phosphorylated complex exhibits a denser interaction pattern, suggesting that phosphorylation promotes additional contacts. Specifically, phosphoserine at position 4 (SEP4) forms interactions with Lys1 and with Gln7 within the peptide, and it also forms a phosphorylation-dependent contact with Asn77 of the HLA molecule. These observations indicate that phosphorylation remodels the local non-bonded interaction network around the modified site by introducing new contacts. In subsequent analyses, we found that the newly formed intra-peptide interactions are stronger than the newly formed peptide–HLA interactions, consistent with a more rigid peptide conformation as described in the following section. This increased rigidity may facilitate stable binding within the HLA groove.

**Figure 2.**
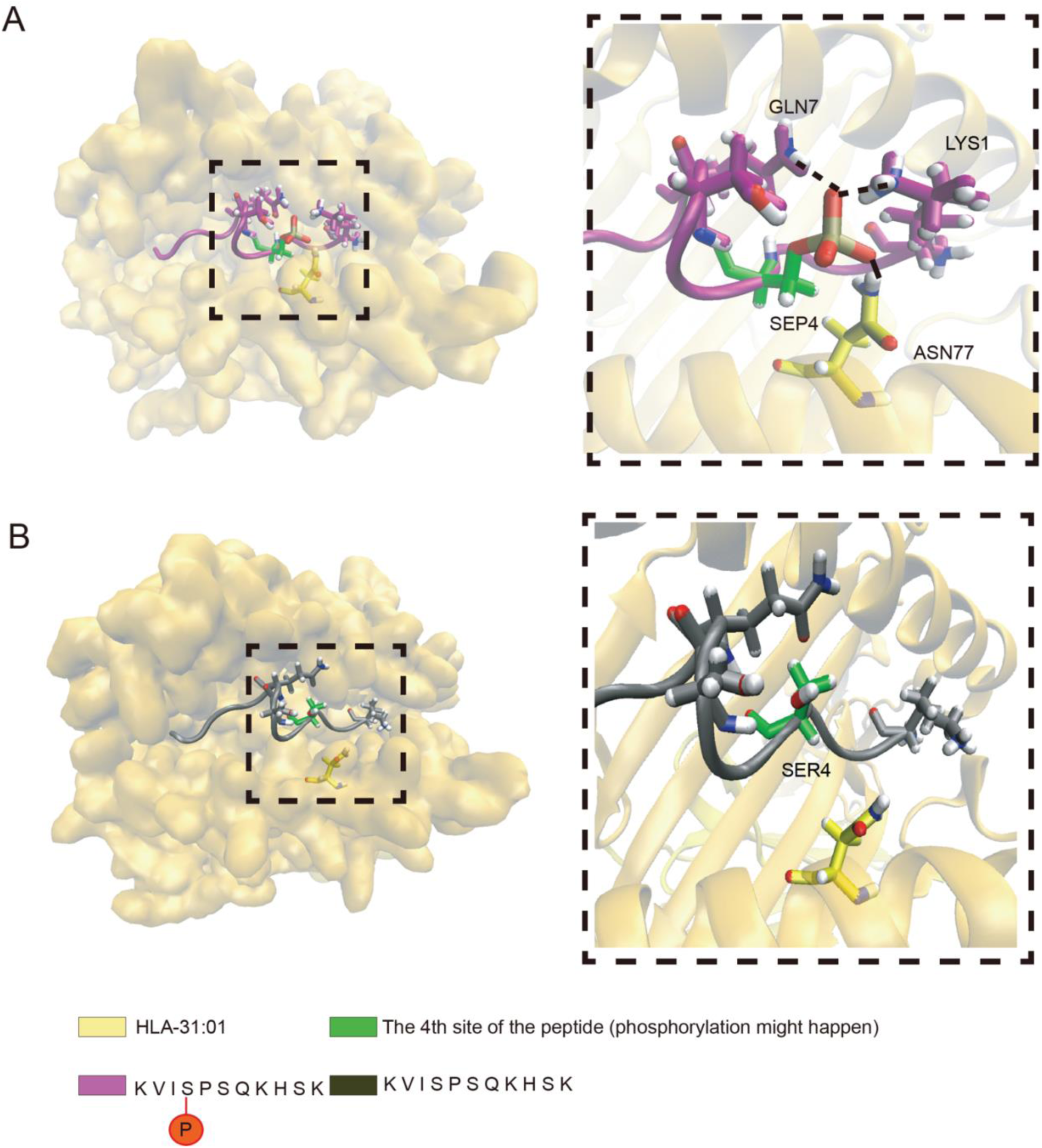
Overall (left) and zoomed-in (right) views of the phosphorylated (A) and non-phosphorylated (B) pHLA complexes. Non-bonded interactions are shown as dashed lines, and interacting residues are labeled.

Beyond the local changes around the phosphorylated site, phosphorylation also reshapes interaction networks across the pHLA complex. As shown in Figure 3, peptide–peptide, peptide–HLA, and HLA–HLA interactions differ substantially between the phosphorylated and non-phosphorylated systems, with some of the largest changes occurring at residues distant from the modified position. To visualize these effects, we present an interaction-difference map in Figure 3D, in which the pHLA structure is colored by the change in interaction strength for each residue. The calculation is defined in Equation 1.

**Figure 3.**
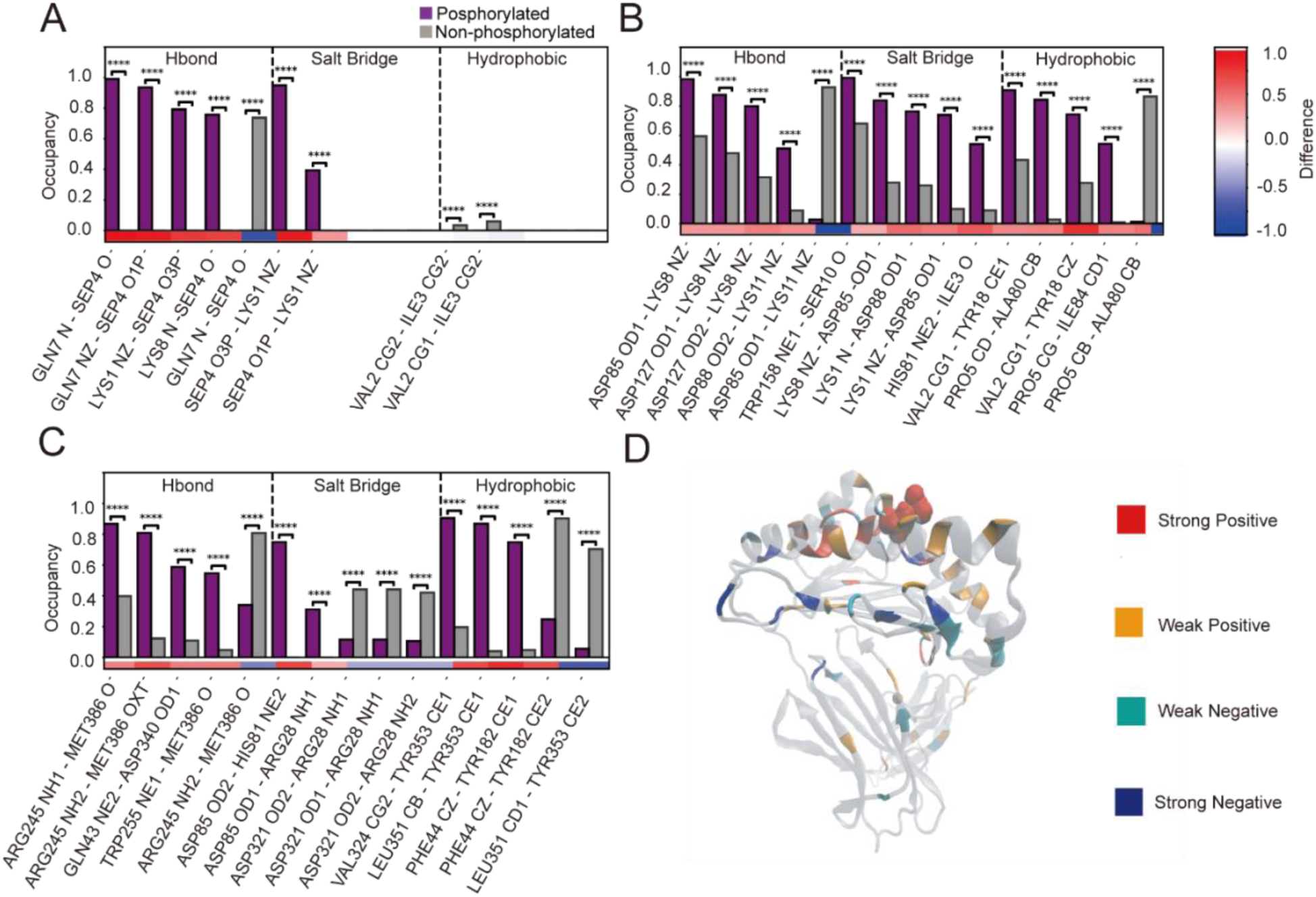
Differences in non-bonded interactions between the phosphorylated and non-phosphorylated systems for peptide–peptide (A), peptide–HLA (B), and HLA–HLA (C) contacts. (D) Residue-level changes in global interaction strength induced by phosphorylation. Strong positive indicates a difference > 0.5; weak positive, 0.2–0.5; weak negative, −0.5 to −0.2; and strong negative, < −0.5. The phosphorylated site is shown in CPK representation.

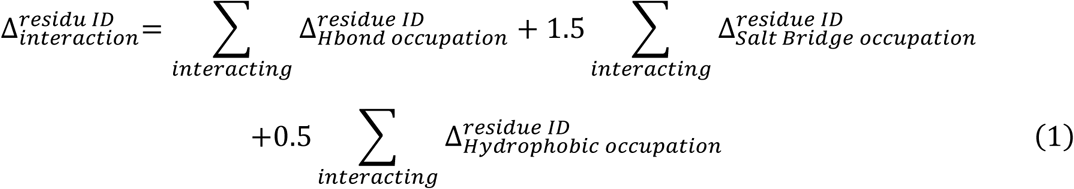

In Equation 1, we assign weights of 1.0 to hydrogen bonds, 1.5 to salt bridges, and 0.5 to hydrophobic interactions, reflecting their typical relative strengths: salt bridges are generally stronger than hydrogen bonds, whereas hydrophobic contacts are weaker on a per-contact basis [39,40]. With these weights, Equation 1 provides a compact approximation of phosphorylation-induced changes in residue-level interaction strength across the network. sNotably, many residues exhibiting large interaction changes are distant from the phosphorylated site, indicating that phosphorylation can propagate global rearrangements in per-residue interaction patterns.

### Phosphorylation enhances pHLA binding, whereas protonation counteracts this effect

Binding affinity is a central consideration in peptide–HLA studies. Here, we estimated peptide–HLA binding affinity using binding free-energy calculations with both the generalized Born (GB) and Poisson–Boltzmann (PB) implicit-solvent models, since the binding free-energy change provides an informative proxy for relative affinity. The results are shown in Figure 4. Although the GB and PB calculations differ in absolute magnitude—reflecting methodological and approximation differences between the two approaches [41]—they yield highly consistent relative trends. Both methods therefore support the same conclusion about how phosphorylation alters peptide–HLA binding affinity. Notably, for the non-phosphorylated peptide, both NetMHCpan [18] and MHCflurry [19] predict weak binding (NetMHCpan: Score_EL=0.461; MHCflurry: mhcflurry_presentation_score=0.525), consistent with the relatively small binding free-energy change observed in our calculations.

**Figure 4.**
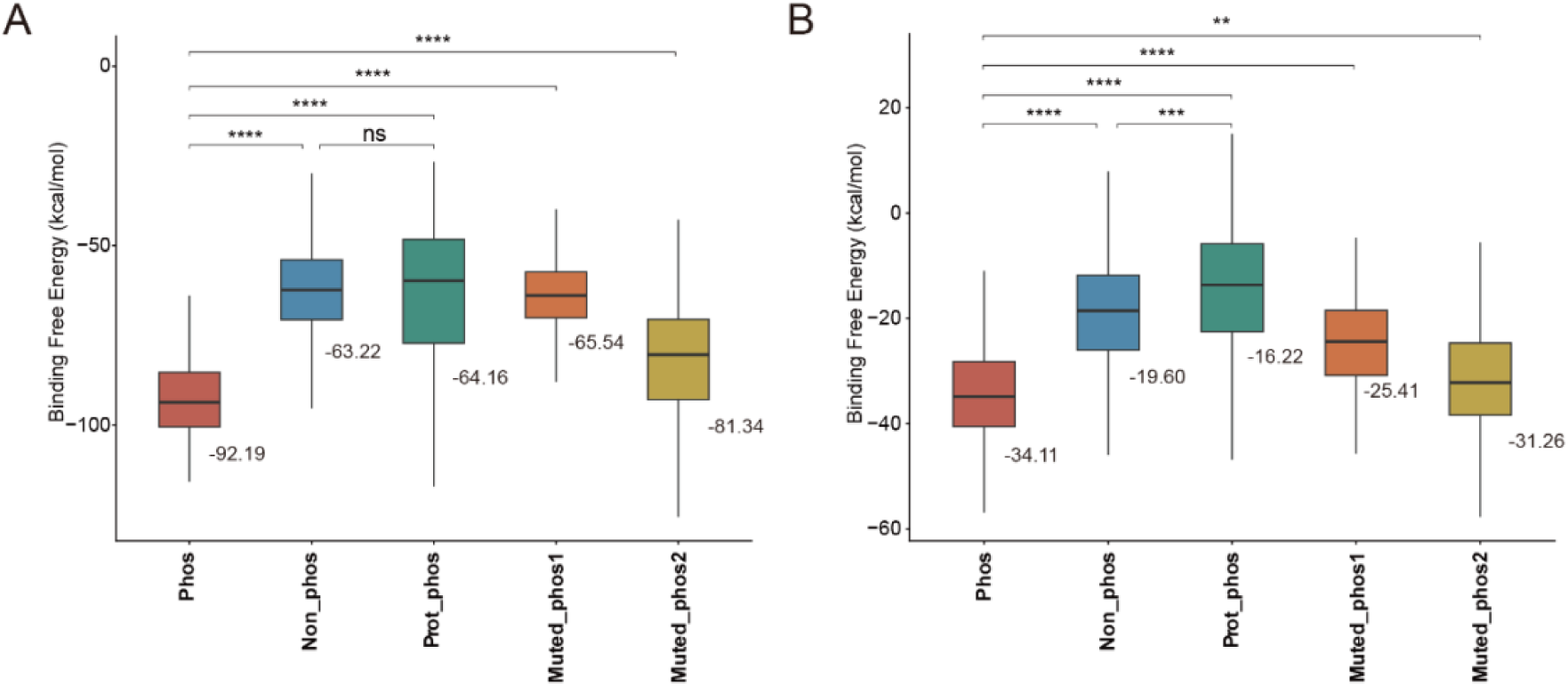
Binding free-energy changes of the pHLA systems estimated using the generalized Born (A) and Poisson–Boltzmann (B) methods. Mean binding free-energy values across simulations are indicated. Phos, Non_phos, Prot_phos, Muted_phos1, and Muted_phos2 denote phosphorylated peptide, non-phosphorylated peptide, the protonated phosphorylated peptide, site 6 phosphorylated peptide, and site 10 phosphorylated peptide, respectively.

These calculations yield several notable observations. First, phosphorylation markedly increases peptide–HLA binding affinity, as reflected by a substantial decrease in binding free energy. Second, in both the GB and PB results, protonation of the phosphorylated peptide reduces binding affinity, with a particularly strong effect in the PB calculations, which use a more detailed electrostatic treatment. Together, these results indicate that although phosphorylation enhances binding, protonation can largely offset this gain.

We also evaluated two additional PTM peptides phosphorylated at alternative sites. Both GB and PB calculations indicate that the correctly phosphorylated peptide has the strongest binding affinity, whereas the incorrectly phosphorylated variants bind more strongly than the non-phosphorylated peptide but weaker than the correctly phosphorylated form. This pattern is consistent with the possibility that the experimentally observed phosphorylation site is preferentially selected because it yields stronger binding to the HLA molecule; however, establishing an evolutionary interpretation would require additional evidence beyond the simulations presented here.

### Phosphorylation reshapes backbone dynamics in the pHLA complex

In addition to binding affinity, phosphorylation may also reshape backbone motions in pHLA complexes. To compare backbone dynamics between the phosphorylated and non-phosphorylated systems, we performed principal component analysis [42] (PCA), a dimensionality-reduction approach that projects conformational fluctuations onto a small number of collective modes. We focused on the first two principal components (PCs), and the results are shown in Figure 5. We carried out four separate PCA analyses for (i) the entire complex, (ii) the peptide alone, (iii) the HLA heavy chain (α chain) alone, and (iv) β2-microglobulin alone. Notably, all projections are reported in a shared PC coordinate system to enable direct comparison across these analyses.

**Figure 5.**
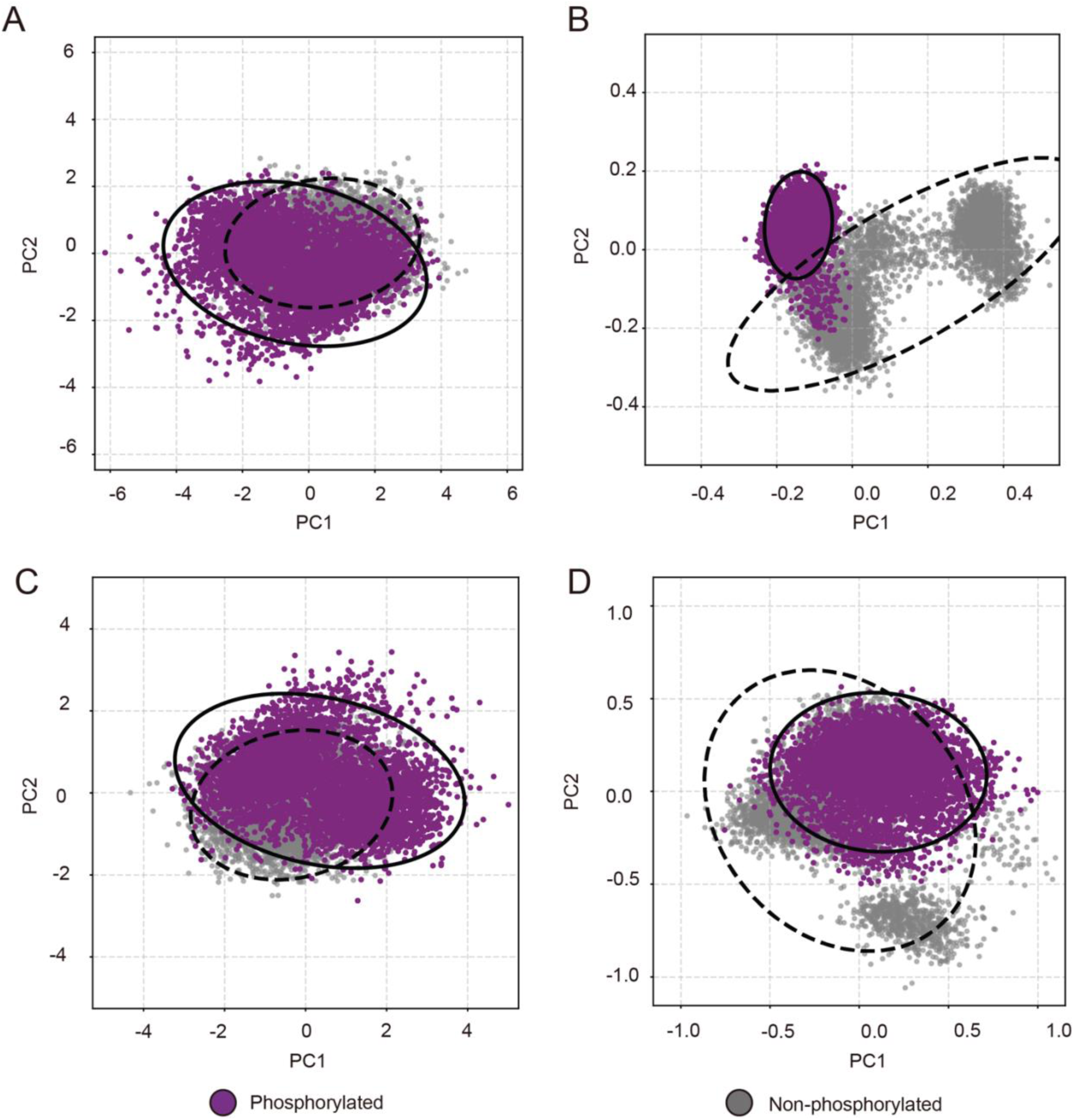
Principal component analysis (PCA) of backbone motions for the phosphorylated and non-phosphorylated pHLA systems, projected onto a shared reference frame. (A) Whole complex, (B) peptide, (C) HLA heavy α chain, and (D) β2-microglobulin. Solid and dashed contours denote the 95% confidence ellipses for the phosphorylated and non-phosphorylated systems, respectively.

From the PCA results, phosphorylation appears to increase the conformational heterogeneity of the overall complex, particularly the HLA heavy chain (α chain), as indicated by a broader distribution of conformations in PC space. In addition, phosphorylation substantially reshapes the dominant backbone motions of both the peptide and β2-microglobulin, consistent with a pronounced change in their dynamical behavior.

To quantify changes in conformational flexibility, we computed ΔS (PCA entropy change) and ΔΔS (difference in entropy change using Equations 2 and 3, respectively. In these calculations, we approximate PCA entropy using the first two PCs [43], and we report both the resulting entropy change and its difference between conditions. Because we define the initial state as a single reference conformation in the PCA map (the weighted average structure of the ensemble), the initial entropy is set to 0. Note that in Equations 2 and 3, R denotes the universal gas constant.

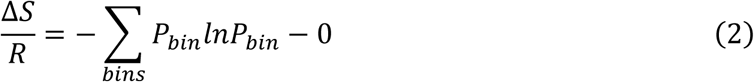

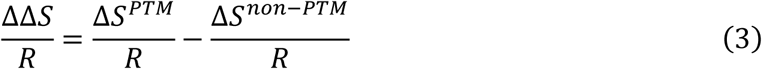

The calculations indicate that phosphorylation yields a ΔΔS/R of −1.242 for the peptide alone and0.303 for the full complex; values for the other components are reported in Supplementary file 2. Together, these results suggest that although phosphorylation strengthens peptide binding to HLA, it increases the overall conformational flexibility of the pHLA system.

We also performed PCA for the protonated phosphorylated system and the non-phosphorylated system (supplementary Figure 5). In these comparisons, the peptide, the HLA heavy chain (α chain), and β2-microglobulin each occupy more dispersed and minimally overlapping regions of PC space, indicating substantial shifts in backbone motions across all three components, as well as in the overall complex. The corresponding entropy analysis results are reported in Supplementary file 2.

## Discussion

In this study, we performed MD simulations and trajectory analyses using a protocol that we first validated for robustness. The simulations indicate that phosphorylation of the serine at position 4 in the ASXL2-derived peptide substantially alters both the binding free energy and the interaction dynamics of the pHLA complex. These findings provide a mechanistic basis that may inform the design of antibodies with specificity for phosphorylated peptide–HLA complexes.

From the binding free-energy calculations and the PCA-based entropy analysis, we infer that the difference in binding affinity between the phosphorylated and non-phosphorylated systems is dominated by the enthalpic contribution. In an NPT ensemble, the free energy change is computed as in Equation 4:

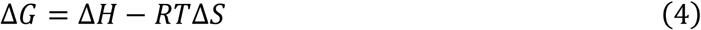

So, the change of the binding free energy change can be evaluated in Eq. 5.

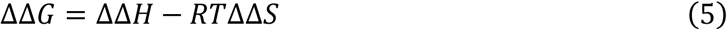

We estimated the enthalpic and entropic contributions to the phosphorylation-induced change in binding free energy by combining (i) the difference in mean PB binding free energies between the phosphorylated and non-phosphorylated systems and (ii) the difference in approximated PCA entropy. This analysis yielded an enthalpic contribution of −14.32 kcal/mol and an entropic contribution of −0.19 kcal/mol, indicating that the change in binding affinity is dominated by enthalpic effects.

The PCA-derived entropy change reflects altered motions of the protein complex, which likely arise from phosphorylation-induced remodeling of interaction dynamics. Notably, peptide-dependent changes in pHLA dynamics have also been reported for non-PTM peptides [26,44]. Consistent with this view, both the non-bonded interaction network and the backbone dynamics shift substantially upon a single phosphorylation on the peptide (Figure 3; Figure 5).

Because the binding-affinity increase is dominated by enthalpic effects and the inferred entropic contribution is small, one might speculate that phosphorylation could enhance immunogenicity by stabilizing peptide–HLA binding. However, such an interpretation is not straightforward because phosphorylation also markedly reshapes pHLA dynamics. As shown in Figure 5, the phosphorylated and non-phosphorylated systems populate distinct regions of conformational space, particularly in the peptide, which is a primary determinant of T cell receptor (TCR) recognition. A phosphorylation-induced shift in peptide conformation could therefore render a TCR optimized for the non-phosphorylated peptide less compatible with the phosphorylated form [45,46]. Under this scenario, immunogenicity could decrease even if peptide–HLA binding becomes stronger, without considering the additional local effects of phosphorylation on charge and sterics.

This interpretation is also consistent with the observation that this phospho-peptide is shared across multiple cancer types and that its modification level increases in many cancers (Figure 1). If phosphorylation were to robustly increase immunogenicity, it would raise the question of why cancer cells bearing this antigen persist. One possibility is immune escape [47] driven, at least in part, by phosphorylation-induced conformational changes in the pHLA complex that alter TCR recognition. In addition, the acidic tumor microenvironment could promote partial protonation of the phosphorylated residue, which our calculations suggest would reduce binding affinity (Figure 4B), potentially further diminishing antigen presentation and immunogenicity.

A practical next step is to develop an antibody with specificity for the phosphorylated form of this ASXL2-derived pHLA complex. As shown in Figure 1, this peptide is observed across multiple cancer types, and phosphorylation is detected specifically in cancer samples. With recent advances in protein design [48]—particularly deep-learning-based approaches [49–51]—it is now timely to ask whether phosphorylation-specific binders can be engineered for this cancer-associated target using state-of-the-art peptide-specific binding protein design strategies [52]. Our results suggest that successful design should account for both the post-translational modification and the accompanying conformational shifts of the pHLA complex. One potential workflow is to design candidate binders against representative structures from the phosphorylated ensemble (for example, a weighted centroid of the MD conformational distribution), then computationally evaluate selectivity by estimating binding free energies for both the phosphorylated and non-phosphorylated complexes, followed by experimental validation. In this context, our study provides a mechanistic basis for in silico design of phosphorylation-specific antibodies and engineered TCRs by clarifying how PTMs can reshape the interaction energetics and dynamics of pHLA systems.

This study has several limitations. First, because the MD simulations are finite in length, ensemble-averaged properties may be sensitive to incomplete sampling and may not fully reflect equilibrium behavior. Second, the chosen force field may introduce inaccuracies, as no force field is universally optimal across all biomolecular systems. Third, PCA-based entropy provides only an approximate proxy for conformational entropy and captures only the collective modes represented in the selected PCs. Fourth, the phosphorylation-induced structural and dynamical changes identified here have not yet been directly validated experimentally. Despite these limitations, our results provide a detailed mechanistic view of how phosphorylation reshapes this antigenic peptide–HLA system and offer hypotheses that can guide future experimental tests and the development of pHLA-targeted cancer immunotherapies.

## Materials and Methods

### Computational materials and data sources

We selected a post-translationally modified (PTM) antigen derived from ASXL2 (phosphorylation) from the caAtlas database [33] and focused on the HLA allele HLA-A*31:01. For comparison, we constructed both phosphorylated and non-phosphorylated pHLA complexes. For the phosphorylated complex, we additionally modeled a protonated state to assess how protonation affects binding affinity and interaction dynamics. We also performed in silico controls in which the peptide was phosphorylated at alternative (incorrect) sites.

The HLA-A*31:01 sequence was obtained from the IPD-IMGT/HLA database [53]. The mature β2-microglobulin sequence was taken from PDB ID: 1AO7 [54]. For computational efficiency, we truncated the intrinsically disordered N-terminal segment and the transmembrane C-terminal region of the HLA heavy (α) chain.

### Data Acquisition and Statistical Analysis

Harmonized CPTAC proteomic and open-access genomic data were obtained via the Proteomic Data Commons (PDC) (https://pdc.cancer.gov/pdc/cptac-pancancer). Raw genomic and transcriptomic data were accessed via the Genomic Data Commons (GDC) Data Portal (https://portal.gdc.cancer.gov; dbGaP Study Accession: phs001287.v16.p6). Controlled datasets hosted in the Cancer Data Service (CDS) were accessed through NCI DAC-approved, dbGaP-compiled whitelists.

### Differential Expression and Pathway Enrichment Analysis

For statistical analysis, differential abundance between tumor and normal tissues was assessed using the Wilcoxon rank-sum test. Pan-cancer significance was evaluated by a signed meta-analysis using Stouffer’s Z-score method to derive a combined Meta P-value. Subsequently, to explore the biological signaling pathways associated with ASXL2 phosphorylation, we performed Gene Set Enrichment Analysis (GSEA). Leveraging the comprehensive proteomic data from the CPTAC dataset, tumor samples were stratified into “High” and “Low” groups based on the median abundance of the specific ASXL2 phosphosite (S156). Differential protein expression analysis was performed between the High and Low phosphorylation groups to generate a ranked gene list (using t-statistics). GSEA was then conducted using the MSigDB Hallmark gene set collection via the clusterProfiler R package.

### Force fields and validation protocol

We used the AMBER ff19SB force field for standard residues [55] and phosaa19SB for phosphoserine and its protonated state [56]. To benchmark whether this force-field setup can distinguish binding from non-binding phosphorylated pHLA systems, we ran short MD simulations in OpenMM 8 using the GB implicit-solvent model [57]. We simulated 50 binding (“positive”) and 50 non-binding (“negative”) phosphorylated pHLA systems at elevated temperature (340 K) for 0.5 ns to enhance conformational sampling within a limited time window. All peptides were constrained to a length of nine amino acids to enable position-wise comparison.

The validation peptides were curated from a large immunopeptidomics dataset of naturally presented phospho-peptides [58], where ligands were assigned to their presenting HLA allele using MixMHCp [59]. The positive set comprised high-confidence phosphorylated ligands from single-allele cell lines. For each positive peptide, we generated a conservative negative control by randomly shuffling the amino-acid sequence while preserving peptide length, modification type, and modified residue identity; the phosphorylation position was allowed to vary. This strategy retains overall composition and PTM state but disrupts the native HLA-binding sequence context. From this pool, we selected 50 matched peptide pairs with unambiguous mapping to HLA-A*31:01 and high confidence in the source dataset.

### Simulation setup and MD protocol

We used AlphaFold3-predicted models as starting structures for geometry optimization [16]. In the AlphaFold3 input, phosphorylated serine was specified using the residue code SEP. Each complex was solvated in a truncated octahedral TIP3P water box [60], with a minimum buffer of 15 Å from the pHLA complex to the box boundary, and 0.15 M NaCl.

All energy minimization, equilibration, and production MD simulations were performed with AMBER 24 [61]. The protocol was as follows. First, we restrained the pHLA complex and ions and minimized only the solvent; we then released restraints and performed a second minimization of the full system. Next, we heated the system from 0 K to 310 K under NVT conditions, followed by 1 ns of NVT equilibration and 1 ns of NPT equilibration. The final coordinates from the NPT stage were used to initialize production simulations. For each pHLA system, we ran a 500 ns production simulation. A 10 Å cutoff was applied for the nonbonded interactions, and the particle mesh Ewald (PME) method [62] was used to calculate the electrostatic interaction with cubic-spline interpolation and a grid spacing of approximately 1 Å.

### Analysis of binding affinity and interaction dynamics

Hydrogen bonds, salt bridges, and hydrophobic contacts were quantified using MDTraj [63]. Backbone Cα RMSD, Cα RMSF, and principal component analysis (PCA) were computed with MDTraj [63]. Binding free energies were estimated using both the generalized Born (GB) and Poisson–Boltzmann (PB) approaches with MMPBSA.py in AMBER 24 [61]. For PB calculations, the dielectric constant was set to 2 for the solute interior and 80 for the solvent. For each system, 200 frames were extracted at evenly spaced time points from the production trajectory for free-energy analysis..

## Resource availability

### Materials availability

This study did not generate new unique reagents.

### Code availability

All codes and scripts we use underlying this manuscript can be accessed from Github at: https://github.com/YiCITI/pASXL2_TumorAntigenMD. The software is distributed under the standard MIT license to facilitate reproducibility and encourage further community development.

## Acknowledgement

This study was supported by the National Natural Science Foundation of China (22304111), the Hundred Talents Program (Category B) of the Chinese Academy of Sciences, and the ‘top Talent’ Program of the Shanghai Advanced Research Institute, Chinese Academy of Sciences (all to X.Y.). Additional support was provided by the Fundamental Research Funds for the Central Universities (YG2025QNA41 to X.Y.). We also acknowledge the computing resources and technical support provided by the Computing Platform (System 9) of the National Facility for Protein Science in Shanghai, Shanghai Advanced Research Institute, Chinese Academy of Sciences.

## Author contributions

X.Y. and J.Z. conceived the study and designed the methodology. X.Y., J.Z., and L.L. performed the formal analysis and wrote the original draft of the manuscript. The investigation was carried out by J.Z., B.C. and L.L. Supervision and funding acquisition were provided by X.Y. All authors reviewed and approved the final manuscript.

## Supplemental Figures

**Supplementary Figure 1.**
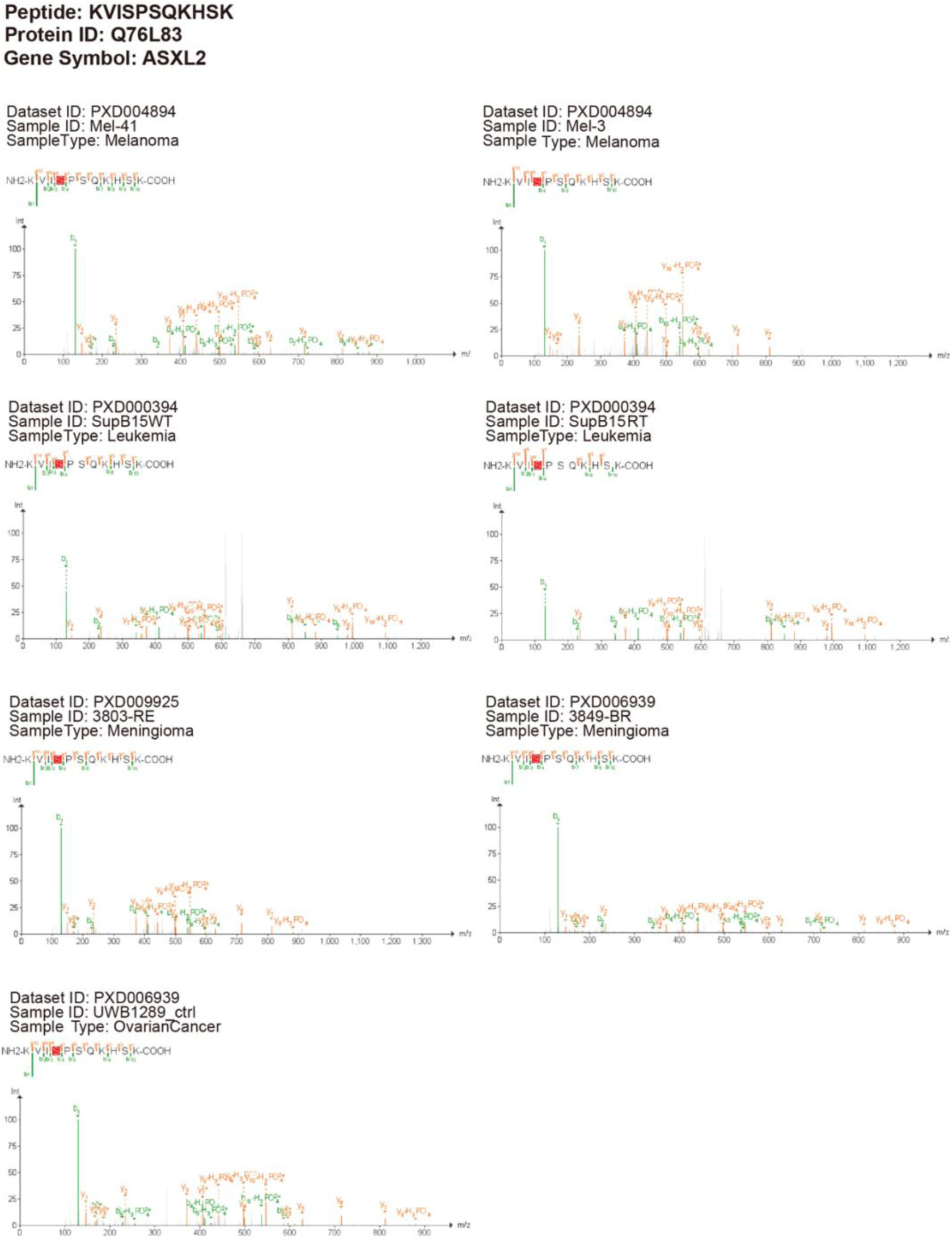
Annotated MS/MS spectra of the ASXL2-derived phospho-peptide retrieved from caAtlas. Representative spectral plots validating the identification of the KVIpSPSQKHSK sequence in the tumor samples shown in Figure 1A. Dataset IDs and specific cancer subtypes are labeled above each spectrum.

**Supplementary Figure 2.**
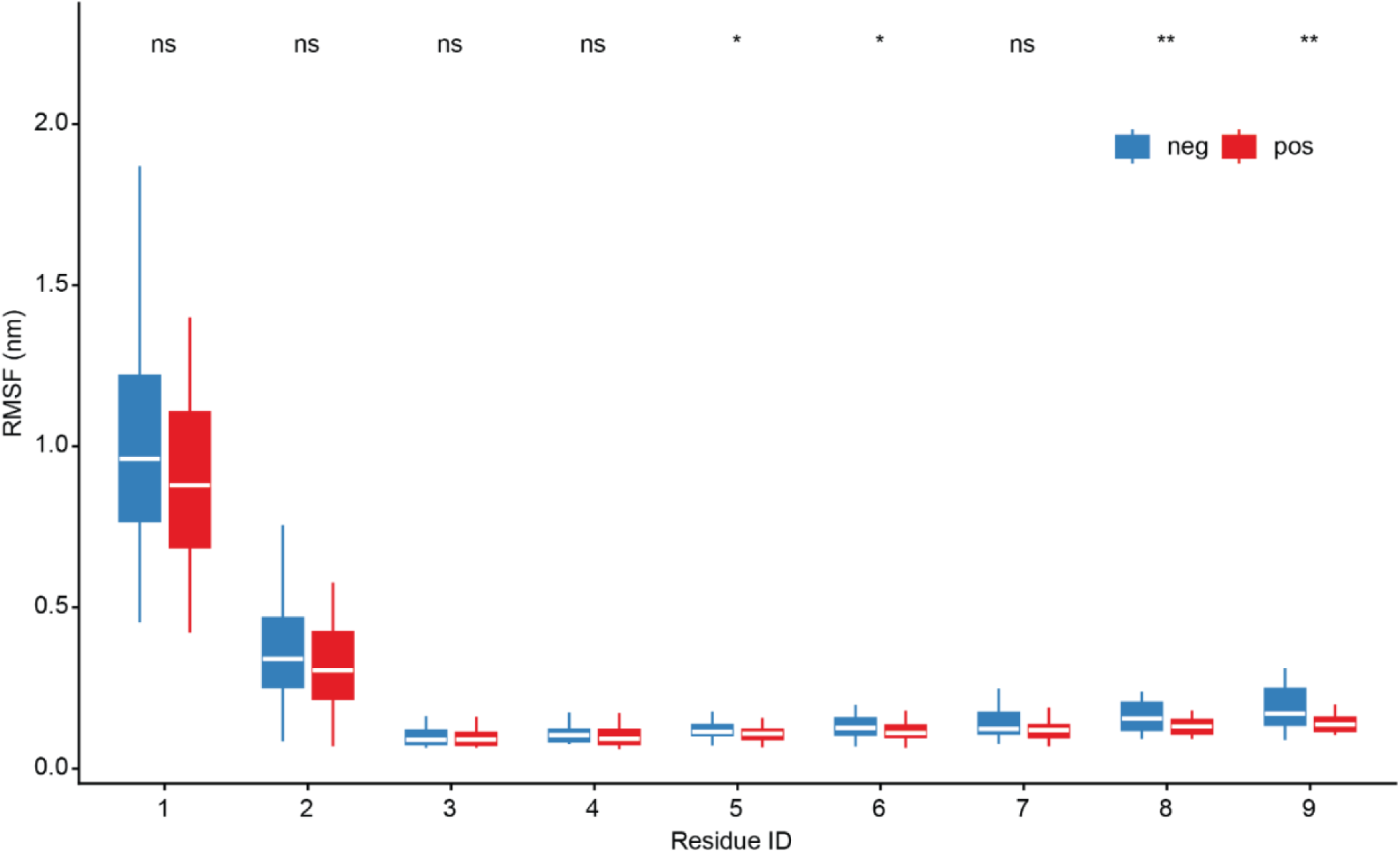
Box plots of per-residue RMSF values for the benchmarking samples.

**Supplementary Figure 3.**
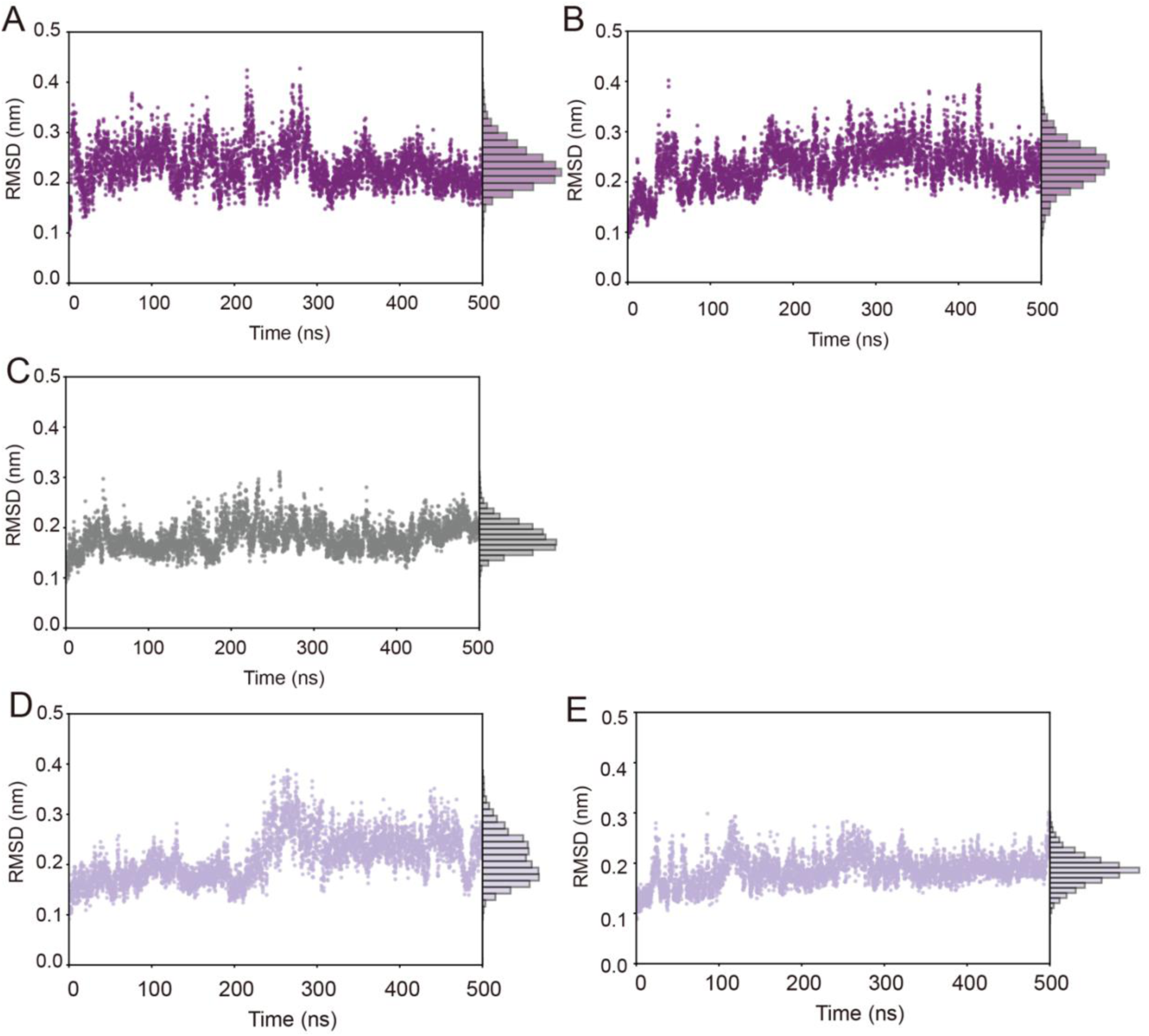
RMSD traces for the production simulations of the phosphorylated system (A), protonated phosphorylated system (B), non-phosphorylated system (C), site 6–phosphorylated system (D), and site 10–phosphorylated system (E).

**Supplementary Figure 4.**
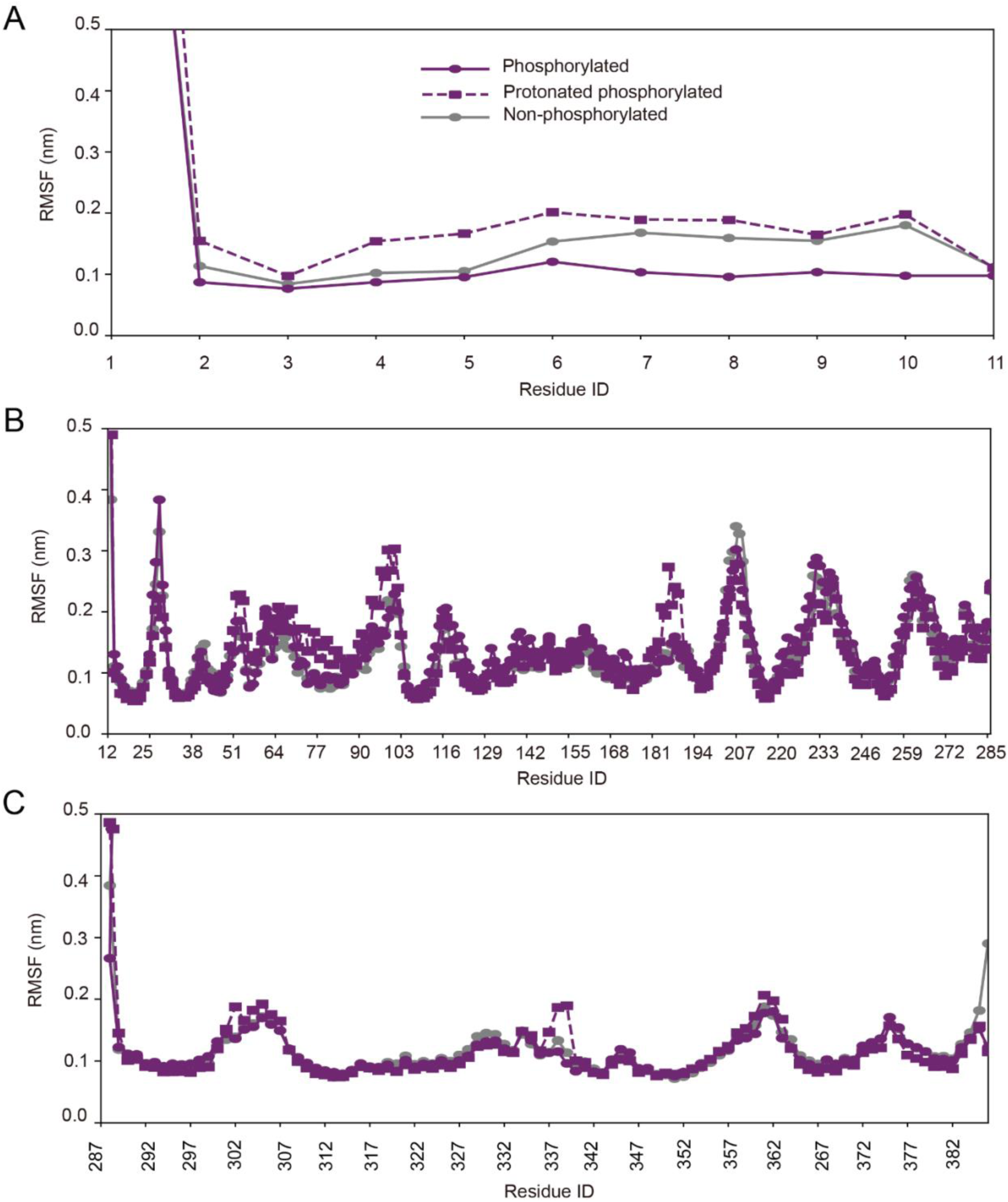
Per-residue RMSF profiles for (A) the peptide, (B) the HLA heavy α chain, and (C) β2-microglobulin.

**Supplementary Figure 5.**
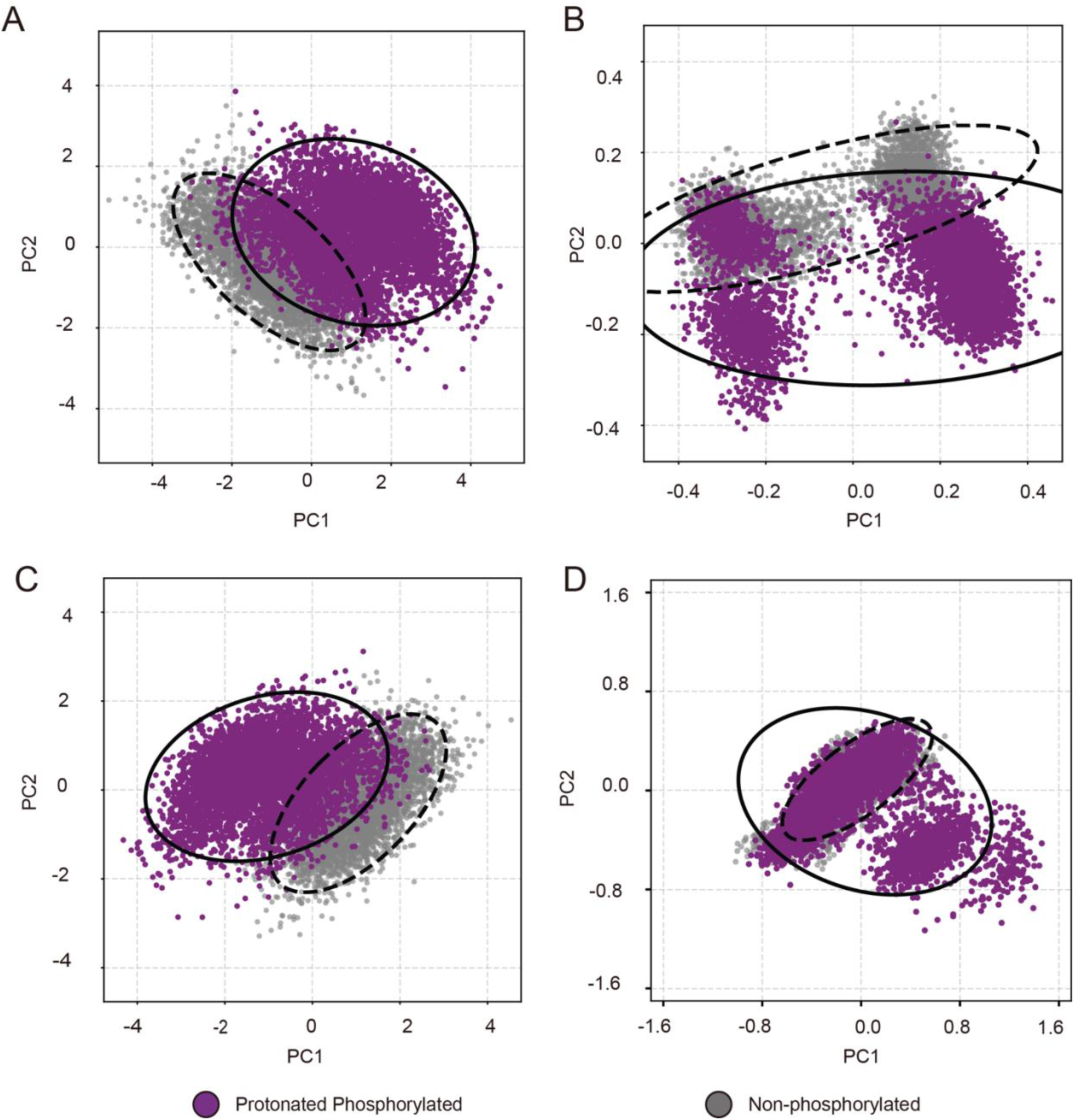
Principal component analysis (PCA) of backbone motions for the protonated phosphorylated and non-phosphorylated pHLA systems, projected onto a shared reference frame. (A) Whole complex, (B) peptide, (C) HLA α chain, and (D) β2-microglobulin. Solid and dashed contours denote the 95% confidence ellipses for the protonated phosphorylated and non-phosphorylated systems, respectively.

## Supplementary file

**1: The samples for force field testing. Supplementary file**

**2: The detailed PCA entropy analysis result. Supplementary file**

**3: The sequences used in this research.**

